# Grasping at the organization of object knowledge: testing different object-related dimensions as organizational principles of ventral temporal cortex

**DOI:** 10.64898/2026.02.10.705013

**Authors:** L. Serriere, G. Argiris, J. Gomes, G.M. Giorjiani, F. Bergström, J. Walbrin, J. Almeida

**Author notes:** Corresponding author: Jorge Almeida, Proaction Lab, Faculty of Psychology and Educational Sciences, University of Coimbra, Portugal.

## Abstract

In our daily lives we encounter a myriad of *things* with which we might need to interact as we navigate our environment. Mental representations of these *things* must be computed and stored in our brains to be manipulated to support cognition. How are such representations organized in the brain? Several proposals have been put forth on what the principles of organization of object information in the brain might be: within ventral temporal cortex – regions thought to support object recognition – possible dimensions include the animacy status of target stimuli, their real size, their texture and material properties, and potentially their graspable status, amongst others. Here we used functional magnetic resonance imaging (fMRI) and multivariate approaches to discriminate patterns of activation for different categories of objects to test the role of these dimensions as organizing principles of object information in the brain. We show that pattern discriminability between different categories of objects does not seem to follow differences in their animacy status in any continuous way. Moreover, graspability of the target stimuli and their haptic texture properties are better predictors of representational content within ventral temporal cortex than animacy and real size. These results are in line with recent studies demonstrating the importance of computational contingencies superimposed by bi-directional functional coupling between parietal regions dedicated to the processing of object manipulation and grasping and ventral temporal regions responsible for object recognition, potentially involving material and texture processing.

## Introduction

Our ability to perceive the world around us and interact with it is supported by the brain’s capacity to abstract away from sensory input, create mental representations of the *things* out there, and match them to stored mental representations. Importantly, the organization of these representations in the brain aids us in distinguishing between different objects and directing us to an appropriate response (e.g., Almeida et al 2023, 2025; Freud et al., 2016; Caramazza & Mahon, 2006). Current data from neuropsychological patients and from neuroimaging experiments with healthy individuals suggests that the Ventral Temporal Cortex (VTC) – an area that putatively subserves object recognition – is organized such that clusters of neurons present response preferences for particular categories of stimuli (e.g., Almeida et al., 2017; Bracci & Op de Beeck, 2023; Chao & Martin, 2000; Epstein & Kanwisher 1998; Kanwisher et al., 1997; Mahon et al., 2007). There is, however, a lack of consensus on the principles that rule this organization – some propose that these response preferences relate to domain-specific processes dedicated to particular types of objects or computations (e.g., Mahon & Caramazza, 2006; Wurm & Caramazza, 2022), others propose overarching dimensions that fully explain this purportedly categorical organization (e.g., Contier et al., 2024; Haxby et al., 2001; Ritchie et al., 2025). In this paper, we test the role of some of these dimensions as principles of organization of the representations in VTC.

Visual object information is thought to be processed along two separate pathways: the dorsal visual pathway and ventral visual pathway (e.g., Goodale & Milner, 1992; Wurm & Caramazza, 2022; Ungerleider et al., 1998; et al; Almeida et al., 2008). These pathways process the incoming visual information to fulfil different computational goals. Particularly, the dorsal visual stream – going from early visual cortex to posterior parietal cortex – is predominantly involved in localizing stimuli in the environment and performing visuomotor computations to support the potential interactions one can have with such stimuli (e.g., Almeida et al., 2010, 2023; Bergström et al., 2021; Culham et al., 2003; Goodale & Milner, 1992); whereas the ventral pathway – going from early visual cortex into lateral and ventral temporal cortex – has been shown to support object recognition and abstract perceptual processing of visual properties of objects such as shape, texture and other surface properties (e.g., e.g., Almeida et al., 2023; Baumgartner and Gegenfurtner, 2016; Cant et al, 2008, 2009; Cant & Goodale, 2007, 2011; Cant & Xu, 2012; Cavina-Pratesi et al., 2010; Grill-Spector et al., 2001; Jacobs et al., 2014; Komatsu & Goda, 2018; Kourtzi & Connor, 2011; Mahon et al., 2007; Prodebarac et al., 2014; Walbrin et al., 2024, Xiao & Liao 2025).

The works of the ventral visual stream – and in particular of the VTC – that support object recognition require, among other things, the ability to generalize and categorize individual exemplars and object types (e.g., my office key; a key) within a general category (a “tool”), differentiating it from other object types within the same category, as well as, potentially object types from other categories (e.g., Grill-Spector & Weiner, 2014; Lee & Almeida, 2021). Interestingly, neuropsychological and neuroimaging data on the processing of objects point to the existence of clusters of neurons that seem, at least at face value, to be dedicated to particular object categories. For instance, some neuropsychological patients presenting with lesions within or around the lateral and ventral temporal cortex – i.e., within the ventral visual stream – seem to show functional deficits that are specific for recognizing exemplars from certain domains, such as faces, manipulable objects, animals, body parts or fruits and vegetables, in the context of spared recognition of exemplars from the other domains (e.g., Caramazza & Shelton, 1998; Garcea et al., 2019; Volfart et al., 2025; Warrington & McCarthy, 1983; Warrington & Shallice 1984; for a review see Capitani et al., 2003). Interestingly, similar category-specific response patterns were obtained in several neuroimaging studies on healthy individuals, where particular regions were found to prefer faces (e.g., Kanwisher et al., 1997; Kanwisher & Novel, 2006), animals (e.g., Chao et al., 1999; Anzellotti et al., 2011), scenes and spatial aspects of the environment (e.g., Aminoff & Durham, 2023; Baldassano et al., 2016; Epstein & Kanwisher 1998; Han & Epstein, 2025), manipulable objects (henceforth tools; e.g., Almeida et al., 2013; Chao & Martin, 2000; Garcea & Mahon 2014; Mahon et al., 2007), or body parts (e.g., Amaral et al., 2021; Bracci et al., 2015; Peelen & Downing 2005) over other categories. Similar responses preferences have been obtained with intracranial recording in humans (e.g., Bastin et al., 2013; Jacques et al., 2013) as well as in experiments with non-human primates (e.g., Bao & Tsao, 2018).

Many views have emerged regarding how to best describe and understand this organization. Some have described it as a sharp functional division reflecting true domain distinctions (e.g., Amaral et al., 2021; Caramazza & Shelton, 1998; Epstein & Kanwisher, 1998; Kanwisher et al., 1997; Kiani et al., 2007; Mahon et al., 2007; Mahon & Caramazza, 2009; Peelen & Downing 2005; Tsao et al., 2008). These domain distinctions underline the evolutionary importance of being able to quickly and reliably distinguish between different entities and perform particular computations for our chances of survival (e.g., Caramazza & Shelton, 1998). These domains can be conceived as first order principles of organization of neural representations (Mahon & Caramazza, 2009), that can then support the kind of multidimensionality that has been shown recently to underlie object processing (Almeida et al., 2023, 2025; Fernandino et al., 2021; Hebart et al., 2023; Huth et al., 2012; Martin, 2016; Valério et al., 2025, 2026), and that are potentially implemented in VTC via connectivity constraints from distal but highly connected regions elsewhere in the brain that code for the same domain (Almeida et al., 2013; Amaral et al., 2022; Bouhali et al., 2014; Bracci & Peelen, 2013; Chen et al., 2017; Garcea et al., 2018, 2019; Hutchinson et al., 2014; Hutchinson & Gallivan, 2018; Kristensen et al., 2018; Lee et al., 2019; Mahon, 2022; Mahon & Caramazza, 2011; Mahon et al., 2013; Osher et al., 2016; Riesenhuber, 2007; Ruttorf et al., 2019; Saygin et al., 2016; Stevens et al., 2015; Walbrin et al, 2024; Walbrin & Almeida, 2021).

Others have proposed that the purportedly categorical nature of the organization of representations in VTC is a consequence of more domain-general organizational principles that are at play within VTC. For instance, some of the regions presenting category-preferences also present particular visual field biases (e.g., face regions within the fusiform gyrus overlap with regions known to prefer foveal stimulation), potentially suggesting that these category-preferring responses (e.g., face-preferring responses) are somehow related with the tendency to foveate particular stimuli (e.g., a face) when processing it (e.g., Levy et al., 2001; Arcaro & Livingstone, 2024). Moreover, some of the category response preferences seem also to overlap with potential differences in how parts of the VTC are tuned to real-object size (Konkle & Caramazza, 2013), potentially to facilitate interactions with objects in the real world and determining object-specific physical constraints that are size-dependent (Konkle & Oliva, 2012).

Alternatively, it has also been proposed that the organization of object representations in VTC is highly distributed. This idea gained further traction with the rise of analytical tools that focused on the patterns of activation across multiple voxels rather than localized activation within a voxel (i.e., multivariate pattern analysis; MVPA; e.g., Haxby & Connolly, 2014; Haynes, 2015; Wang et al., 2013). For instance, Connolly and colleagues (Connolly et al., 2012) used Representational Similarity Analysis (RSA; Kriegeskorte et al., 2008) to show that the patterns of activation along the Lateral Occipital Cortex could be explained by subjective (general) similarity ratings over the stimuli presented. Moreover, O’Toole and colleagues (O’Toole et al., 2005) found that using data from the whole VTC allowed for reliable detection of the kinds of stimuli presented to participants. Importantly, they did so significantly better than when using data from voxels maximally activated for a given stimulus (i.e., category-preferring regions). These results paved the way for the idea that object representations are graded and distributed along particular dimensions.

In line with this, Sha and colleagues (Sha et al., 2015; see also Connoly et al., 2012; Proklova et al., 2016; Ritchie et al., 2021; Ritchie & Op de Beeck, 2019; Thorat et al., 2019) investigated the importance of one possible overarching and distributed dimension – animacy – in determining the functional organization of the VTC and the representational nature of its patterns of activation. They used Principal Component Analysis (PCA) to uncover the latent dimensions organizing neural representational spaces within VTC. They showed that the primary dimension seemed to follow (continuously) the animacy status of each object type. Overall, these reports strongly suggest that animacy – as an overarching dimension that continuously organizes all exemplars of the tested categories – can explain the response preferences observed in VTC.

While the reliance on such animacy continuum as the dominant determinant of the VTC’s functional organization is compelling and follows the relatively consensual hypothesis of the importance of the animate/inanimate divided in understanding mental representation of objects (e.g., Caramazza & Shelton, 1998; Konkle & Caramazza, 2013; Kriegeskorte et al., 2008), it may also face some challenges. Recent work has been demonstrating that objects can be described along multiple dimensions that are important for an object’s representation and many of which are coded within VTC (e.g., Almeida et al., 2023, 2025; Fernandino et al., 2021; Hebart et al., 2020; Huth et al., 2012, Valério et al., 2025, 2026). Moreover, visual properties unrelated to animacy, such as shape, texture, color and material have been shown to be processed along the lateral and ventral temporal aspects of the temporal and occipito-temporal cortex and thus may contribute to the representational nature of VTC. In particular, object texture and material properties are thought to be partly computed along the more medial aspects of the ventral stream (e.g., Almeida et al., 2023; Baumgartner and Gegenfurtner, 2016; Cant et al, 2008, 2009; Cant & Goodale, 2007, 2011; Cant & Xu, 2012; Cavina-Pratesi et al., 2010; Grill-Spector et al., 2001; Jacobs et al., 2014; Komatsu & Goda, 2018; Kourtzi & Connor, 2011; Prodebarac et al., 2014; Xiao & Liao 2025). These data suggest other possible dimensions as central for explaining the organization of object representations in VTC.

Furthermore, the organization of representations within VTC may also be dependent on information not typically associated with ventral stream processing – namely grasping and other action-related properties of objects (Almeida et al., 2008, 2010, 2013, 2018; Amaral et al., 2021; Chen et al., 2017; Cortinovis et al., 2025; Garcea et al., 2018, 2019; Hussain et al., 2024; Kristensen et al., 2016; Lee et al., 2019; Mahon, 2013, 2022; Ruttorf et al., 2019; Walbrin et al, 2024; Walbrin & Almeida 2021; for a review see Mahon & Almeida, 2024) – via connectivity constraints and processing contingencies (Mahon & Almeida, 2024). While Mahon et al. (2007) had already demonstrated that part of the ventral stream activates selectively according to motor-relevant properties of objects, several data show important bidirectional connectivity between ventral and parietal areas, with parietal areas supporting ventral stream activations responsible for object recognition (e.g., Almeida et al., 2013; Amaral et al., 2022; Chen et al., 2017; Garcea et al., 2018, 2019; Hutchinson et al., 2014; Hutchinson & Gallivan, 2018 Kanwisher, 2016; Kristensen et al., 2018; Lee et al., 2019; Mahon, 2022; Mahon & Caramazza, 2011; Mahon et al., 2013; Osher et al., 2016; Riesenhuber, 2007; Ruttorf et al., 2019; Saygin et al., 2016; Stevens et al., 2015; Walbrin et al, 2024; Walbrin & Almeida, 2021; for a review see Mahon & Almeida, 2024). Consequently, an object’s graspable status (Almeida et al, 2010; Almeida & Mahon, 2024) may also challenge the dominance of animacy as an organizing principle of the neural representations in VTC.

Thus, we initially set out to explore the role of animacy (and the animacy continuum), as a major contender for explaining response preferences in VTC. To do so, we presented stimuli from a set of categories: some of which clearly fall within those typically used to test both category-specificity and animacy representations in VTC – the categories of faces, animals (represented here by the subordinate category of mammals) and tools; and some of which provide, in our view, leverage to test animacy as a major organizational principle of neural representations – the categories of fruits and vegetables and of insects. The category of fruits and vegetables is a central category in the processing of high-level information and in the understanding of neural and mental object representation. This is so because 1) neuropsychological patients can be impaired specifically at recognizing exemplars from this category in the context of spared recognition of exemplars from other categories (Laiacona & Capitani, 2001; Samson & Pillon., 2003); 2) exemplars from this category have been shown to drive response preferences in particular regions of the VTC (Avery et al., 2022; 2025; Bannert & Bartels, 2022; Jain et al., 2023; Khosla et al., 2022; Ritchie & Op de Beeck, 2019); and, perhaps more importantly, 3) they pose a challenge to many of the important views in the organization of conceptual knowledge, in that they are living things that are nevertheless inanimate. The category of insects also provides an opportunity to address these issues as it is certainly a category of animate entities, and especially so when compared to fruits and vegetables. Thus, if animacy is the major organizing principle of response preferences in VTC, then insects will necessarily be more similar to other animate things like faces (and less similar to inanimate things like tools) than fruits and vegetables.

To test this, we first looked at neural discriminability between the responses to animals, insects and fruits and vegetables against tool and face stimuli. Specifically, we used MVPA and linear Support Vector Machine (SVM) algorithms to train classifiers to distinguish the patterns of activation of each of these stimulus categories binarily against the two baseline animate/inanimate categories (i.e., faces and tools). We show that unlike what would be predicted by the animacy hypothesis, fruits and vegetables and insects are equally hard to discriminate from tool stimuli – if anything discriminating fruits and vegetables from tools is numerically easier – and are equally easier to discriminate against face stimuli – if anything discriminating insects from faces is numerically easier.

We then focused on testing a larger set of dimensions whose computations are thought to be associated with the processing within VTC – e.g., animacy, haptic texture, visual texture, shape, real size, and graspability – and that are potential organizing principles of the representations computed and stored within VTC. Using RSA, we compared representational content within the neural activations for the different categories to multiple behavioral models independently obtained that represent those different dimensions. Given previous work showing the importance of grasp and action-relevant information and an object’s real-size in visual object processing and recognition, our focus is on the models subserving these processes: models related with texture that has the potential of informing how we grasp and manipulate and object (haptic texture) and the graspable status of an object, as well as on an objects real-size, along with an object’s animacy status. Our findings show that the models that best explained the representations within VTC are indeed those related to object-directed action – haptic texture and graspability. We discuss our results in the context of the organization of object representations in VTC.

## Methods

### Participants

16 individuals (13 women; mean age ± standard deviation = 20.44 ± 3.24) from the community of the University of Coimbra participated in the main (fMRI) experiment. Moreover, 108 individuals (14 men; mean age ± standard deviation = 20,8 ± 3,7) were recruited for the behavioral experiments to obtain dimensional models for testing the representational content within VTC. All participants were right-handed, had normal or corrected-to-normal vision, and were naive as to the experimental manipulations. The experiment was approved by the ethics committee of the Faculty of Psychology and Educational Sciences of the University of Coimbra and followed all ethical guidelines. Moreover, participants provided written informed consent, where they were informed of the requirements to participate in an fMRI experiment (e.g., no metal). Participants were compensated for their time by receiving either course credit on a major psychology course or financial compensation.

### Materials and Procedure

#### Stimuli

Images of objects belonging to one of five categories were used: animals (mammals), fruits and vegetables (F&V), (familiar) faces, tools, and insects. We selected twelve types for each category, with four exemplars per object type (total of 240 images) The object types per category were the following: Faces (*Halle Berry, Sandra Bulllock, Angelina Jolie, Natalie Portman, Julia Roberts, Kate Winslet, Alec Baldwin, Tony Blair, George Bush, Tom Cruise, Matt Damon, Robert De Niro*); Animals (*bear*, *cat*, *cow*, *deer*, *dog*, *elephant*, *giraffe*, *horse*, *lion*, *monkey*, *pig rabbit*); Fruits and Vegetables (*apple, banana, broccoli, cabbage, cherry, eggplant, mushroom, pepper, pineapple, pomegranate, strawberry, tomato*); Insects (*ant, bee, beetle, butterfly, caterpillar, dragonfly, firefly, fly, grasshopper, ladybug, moth, wood louse*); and Tools (*Corkscrew, fork, hammer, knife, pan, pliers, razor, saw, scissors, screwdriver, shovel, teapot*). Images were collected from the World Wide Web or taken from internal image databases or the FRIDa database (Foroni et al., 2013). Pictures were then converted to grayscale, downsized to 400 x 400 pixels, and presented on a gray background. Stimuli were matched for brightness between categories (F (4, 235) = .415, p = .8).

#### fMRI Procedure

Participants underwent a standard multicategory block-design task (e.g., Almeida et al., 2013; Fintzi & Mahon, 2014; Lee et al., 2019), where they were asked to passively view a series of images of objects from different categories (see Figure 1). In each run, two miniblocks for each category were presented. Each miniblock consisted of 24 images of one category that were presented sequentially for 500 milliseconds each (total of 12 s). Right after each miniblock, a 12s period of fixation was presented. Each run started and ended with 16 s of fixation to allow signal stabilization and thus lasted approximately 4 minutes 32 seconds. Participants went through 10 runs. Stimuli were backprojected on an LCD monitor and subtended approximately 10° of the visual angle. Stimulus presentation was controlled using Psychtoolbox in MATLAB (Brainard, 1997; Kleiner et al, 2007).

**Figure 1.**
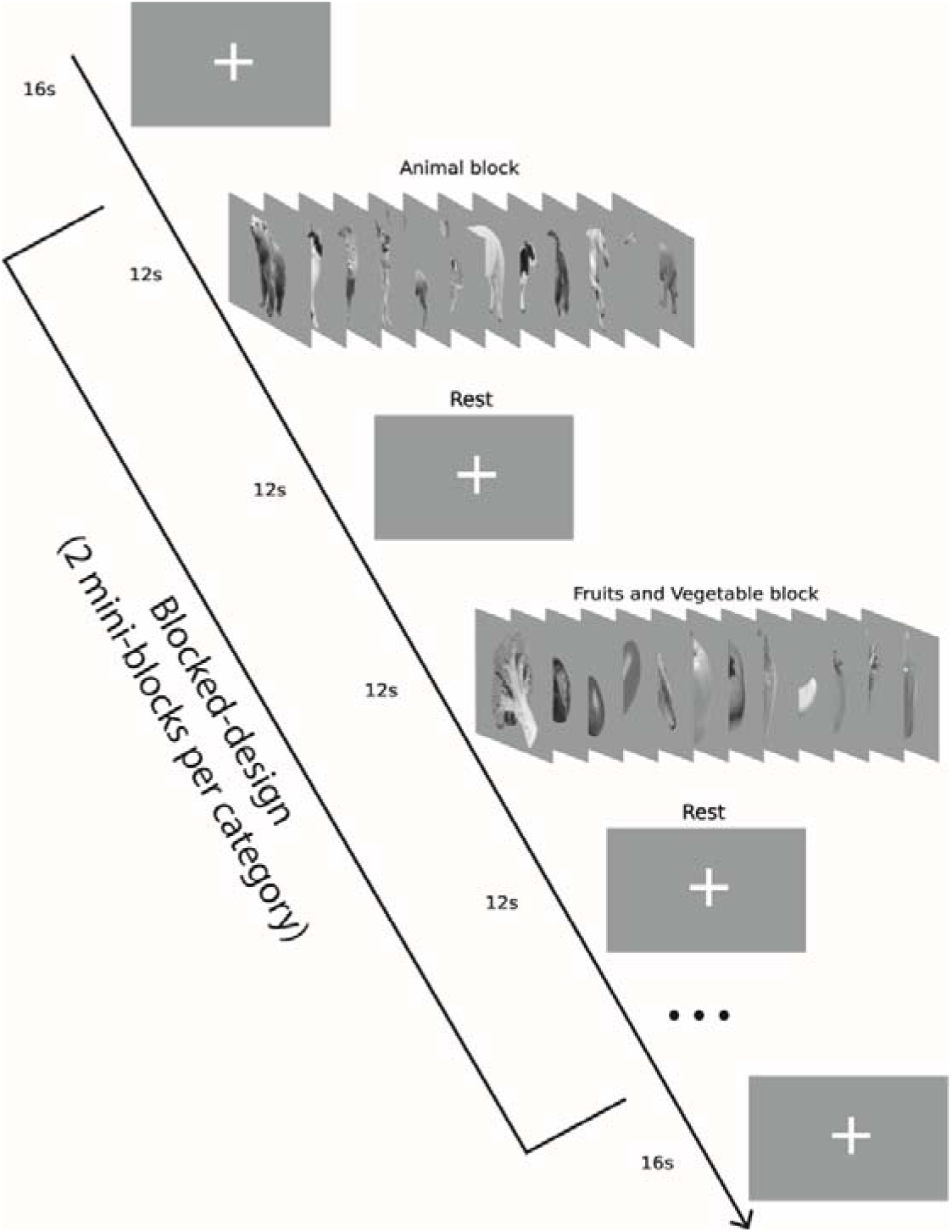
fMRI experimental design. A block design was used to investigate the functional processing of different object categories. Each run started with a 16s fixation cross, followed by 10 mini-blocks of stimulus presentation. Each mini-block consisted of 24 images presented for 500ms each, followed by a 12s fixation cross mini-block. A final fixation cross was presented at the center of the screen for 16s at the end of the run. The experiment consisted of 10 runs, lasting approximately 50 minutes.

#### Behavioral Procedures

To understand how the different categories are perceived along several object-related dimensions of interest, we asked participants to perform one-dimensional arrangements of the different stimuli according to each of the target dimensions. All objects were rated in 18 different measures. Due to the long task duration (approximately 10 minutes for each rating dimension), these measures were collected from 4 different groups of participants. For most of the dimensions tested, participants were given two anchors and were asked to place the different images (those also used in the fMRI experiment) according to their “value” on that dimension. The dimensions tested included aspects related to animacy, such as the degree of self-awareness (anchors: 0% and 100%), the ability of self-motion (anchors: 0% and 100%), and the level of cognitive functioning (anchors: 0% and 100%); aspects related to the ontological status of an object, namely artificiality of the object (anchors: artificial and natural), and whether it is handmade or naturally occurring (anchors: handmade and natural); aspects related to texture and material properties of the stimuli such as smoothness (anchors: smooth and rough), softness (anchors: soft and hard), furriness (anchors: smooth and furry), temperature (anchors: cold and warm), texture complexity (anchors: simple texture and complex texture), glossiness (anchors: mate and glossy), opaqueness (anchors: opaque and transparent) and color (anchors: colorless and colorful); an aspect related to object manipulation, grasping (easy to grasp and hard to grasp) which was determined as the ease with which one could hold the whole object with one hand; and aspects related to the shape of an object, such as real size (anchors: small and big). We also collected other shape-related dimensions, such as elongation (anchors: elongated and concentric), edge (anchors: sharp edge and curvilinear), general visual similarity, form similarity, among the different objects. The latter two dimensions were anchorless, we used the inverse MDS arena technique (Kriegeskorte & Mur, 2012), whereby participants have to drag images of the stimuli presented on a screen to a centrally-presented circular arena, positioning them according to how similar those objects are to one another. The placing of each stimulus is then used for calculating similarity-based distance values.

Overall, data from 6 participants were excluded due to non-completeness of the task, or data corruption. To determine how the categories ranked on each measure, the rating of the exemplars within each category was averaged. Different dimensions were then combined to create meaningful behavioral models. Specifically, we created the following models: Animacy (the simple average of the dimensions Self-Awareness, Self-motion, and Cognitive Functions), Haptic Texture (the simple average of the dimensions smoothness, softness, furriness, and temperature), Visual Texture (the simple average of the dimensions texture complexity, glossiness and opaqueness), Real Size, Visual Shape (the simple average of the dimensions general visual similarity and form similarity), and Artificiality (the simple average of the dimensions artificiality and handmadeness). See supplementary table S1 for the scores of each category for each model.

### Analysis

#### MRI acquisition

Scanning was performed with a Siemens MAGNETOM Prisma-fit 3T MRI Scanner (Siemens Healthineers) with a 32-channel head coil at the University of Coimbra (Portugal; BIN - National Brain Imaging Network). Functional images were acquired with the following parameters: T2* weighted (single-shot/GRAPPA) echo-planar imaging pulse sequence, repetition time (TR) = 2000 ms, echo time (TE) = 22 ms, and voxel size of 2.3 mm^3^, field of view (FOV) = 220 mm; acquisition matrix = 256 x 256 (40 descending interleaved even-odd slices). Structural T1-weighted images were obtained using a magnetization prepared rapid gradient echo (MPRAGE) sequence with the following parameters: TR=1900, TE=2.3 ms, acquisition matrix = 256 × 256, and voxel size of 0.9 mm^3^ (singleshot ascending slice sequence). Foam padding was used to reduce head movement within the coil.

### Pre-processing of (f)MRI data

#### Anatomical

Data was preprocessed using fMRIprep version 20.2.0 (Esteban et al., 2018). The T1w image was corrected for intensity non-uniformity (INU) with N4BiasFieldCorrection (Tustison et al. 2010), distributed with ANTs 2.5.3 (Avants et al. 2008, RRID:SCR_004757), and used as T1w-reference throughout the workflow. The T1w-reference was then skull-stripped with a Nipype implementation of the antsBrainExtraction.sh workflow (from ANTs), using OASIS30ANTs as target template. Brain tissue segmentation of cerebrospinal fluid (CSF), white-matter (WM) and gray-matter (GM) was performed on the brain-extracted T1w using fast (FSL (version unknown), RRID:SCR_002823, Zhang, Brady, and Smith 2001). Brain surfaces were reconstructed using recon-all (FreeSurfer 7.3.2, RRID:SCR_001847, Dale, Fischl, and Sereno 1999), and the brain mask estimated previously was refined with a custom variation of the method to reconcile ANTs-derived and FreeSurfer-derived segmentations of the cortical gray-matter of Mindboggle (RRID:SCR_002438, Klein et al. 2017). Volume-based spatial normalization to one standard space (MNI152NLin2009cAsym) was performed through nonlinear registration with antsRegistration (ANTs 2.5.3), using brain-extracted versions of both T1w reference and the T1w template. The following template was selected for spatial normalization and accessed with TemplateFlow (24.2.0, Ciric et al. 2022): ICBM 152 Nonlinear Asymmetrical template version 2009c [Fonov et al. (2009), RRID:SCR_008796; TemplateFlow ID: MNI152NLin2009cAsym].

#### Functional

First, a reference volume was generated, using a custom methodology of fMRIPrep, for use in head motion correction. Head-motion parameters with respect to the BOLD reference (transformation matrices, and six corresponding rotation and translation parameters) are estimated before any spatiotemporal filtering using mcflirt (FSL; Jenkinson et al. 2002). The estimated fieldmap was then aligned with rigid-registration to the target EPI (echo-planar imaging) reference run. The field coefficients were mapped on to the reference EPI using the transform. The BOLD reference was then co-registered to the T1w reference using bbregister (FreeSurfer) which implements boundary-based registration (Greve and Fischl 2009). Co-registration was configured with six degrees of freedom. Several confounding time-series were calculated based on the preprocessed BOLD: framewise displacement (FD), DVARS and three region-wise global signals. FD was computed using two formulations following Power (absolute sum of relative motions, Power et al. (2014)) and Jenkinson (relative root mean square displacement between affines, Jenkinson et al. (2002)). FD and DVARS are calculated for each functional run, both using their implementations in Nipype (following the definitions by Power et al. 2014). The three global signals are extracted within the CSF, the WM, and the whole-brain masks. Additionally, a set of physiological regressors were extracted to allow for component-based noise correction (CompCor, Behzadi et al. 2007).

#### Ventral Temporal Cortex (VTC) ROI

The VTC was defined using Juelich brain’s atlas in SPM’s Anatomy toolbox (Eickhoff et al., 2007). Four cytoarchitecturally defined fusiform gyrus slices per hemisphere are defined in their atlas. Taken together, these areas constitute a large part of the temporal cortex responsible for the processing of visual stimuli (MNI coordinates: Right VTC: 24 < x < 54, -75 < y < -28, -31 < z < -2, 741 voxels; Left VTC; -21 < x < -52, -79 < y < -28, -31 < z < -2, 958 voxels). This template allowed us to explore the functioning of the whole VTC by combining the different fusiform regions detailed in the atlas (FG1 - FG4).

#### Neural discriminability of the different categories of stimuli

An initial univariate analysis was performed to extract the relevant functional activation for our conditions of interest. First and second level analyses were performed using SPM12 (Welcome Trust Centre for Neuroimaging, London, UK) on data smoothed to a 3mm kernel (Hendriks et al., 2017). All parameters were kept to the standard SPM options. A General Linear Model (GLM) was applied to the neural data to detect the blood oxygenation levels dependent (BOLD) on the conditions of interest. 19 nuisance regressors extracted using fMRIprep were used in the GLM analysis. These included 6 motion parameters (translation and rotation for each x, y and z dimensions), as well as their first-order temporal derivatives and squared terms, and the framewise displacement to account for head motion artifacts. These regressors were subtracted from the GLM which allowed for the extraction of beta values most reflective of the true hemodynamic response elicited by the different conditions.

The SVM decoding models were subsequently trained on the beta values extracted from the first level analysis. Specifically, a within-subject leave-one-run-out approach was used to avoid circularity, where the training set was composed of nine runs and the model was tested on the left-out run. This resulted in 10 folds per subject, ensuring each run was used as a testing sample. A z-score normalisation of the beta patterns was additionally computed using the training set data and applied to the test sample for each fold. Binary classifications were performed against the two categories considered to be the extremes of a potential animacy continuum – Faces and Tools. Thus, we trained SVM classifiers to discriminate activations elicited by tools from those elicited by fruits and vegetables, insects, animals and faces, and to discriminate activations elicited by faces from those elicited by fruits and vegetables, insects, animals and tools. The analyses were performed for activation patterns from all the voxels within the VTC ROIs described above. We used the default CoSMoMVPA SVM classifying function (Oosterhof et al., 2016) and computed the significance between classification performances using FDR corrected t-tests with false discovery rate of q<0.05 for all tests within a decoding paradigm (four classifications versus faces, and four classifications versus tools).

#### Representational content within the VTC

To further understand the neural organization of the information in VTC, we used Representational Similarity Analysis (RSA; Kriegeskorte et al., 2008) to explore the representational content of the VTC against the behavioral models obtained. To do so we computed Representational Dissimilarity Matrices (RDMs) for the neural responses using CoSMoMVPA (Oosterhof et al., 2016) of the left and right VTC. This process whitened the noise and averaged the beta value of each voxel per participant for each condition. The pairwise dissimilarity of each category was calculated by initially centering the data by subtracting the grand mean from each value, then calculating the Spearman’s correlation between each category pair. For the behavioral RDMs, the difference of the scores between each stimulus for a given model was first calculated and subsequently averaged per condition. The euclidean distance between each condition was then computed and standardized to be re-ranged between 0 and 2. These can be seen in figure 2. The behavioral RDMs are shown in figure 3.

**Figure 2.**
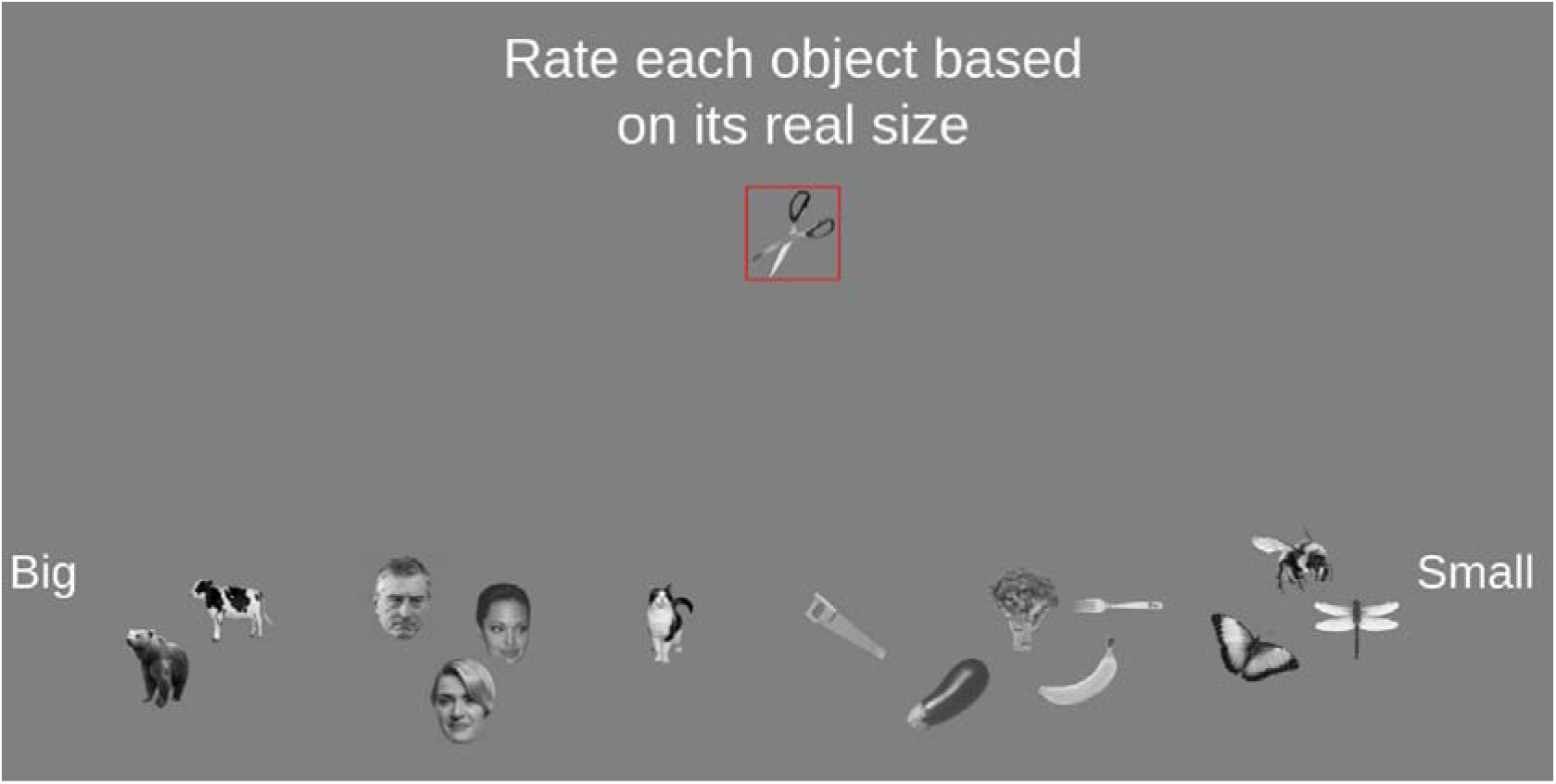
Behavioral experimental design. Participants were tasked with placing all 60 images along a measure’s continuum. This was done one image at a time with new images appearing at the top, outlined in red. Once placed, the stimuli could still be moved until the participant indicated they were satisfied with the overall placement.

**Figure 3.**
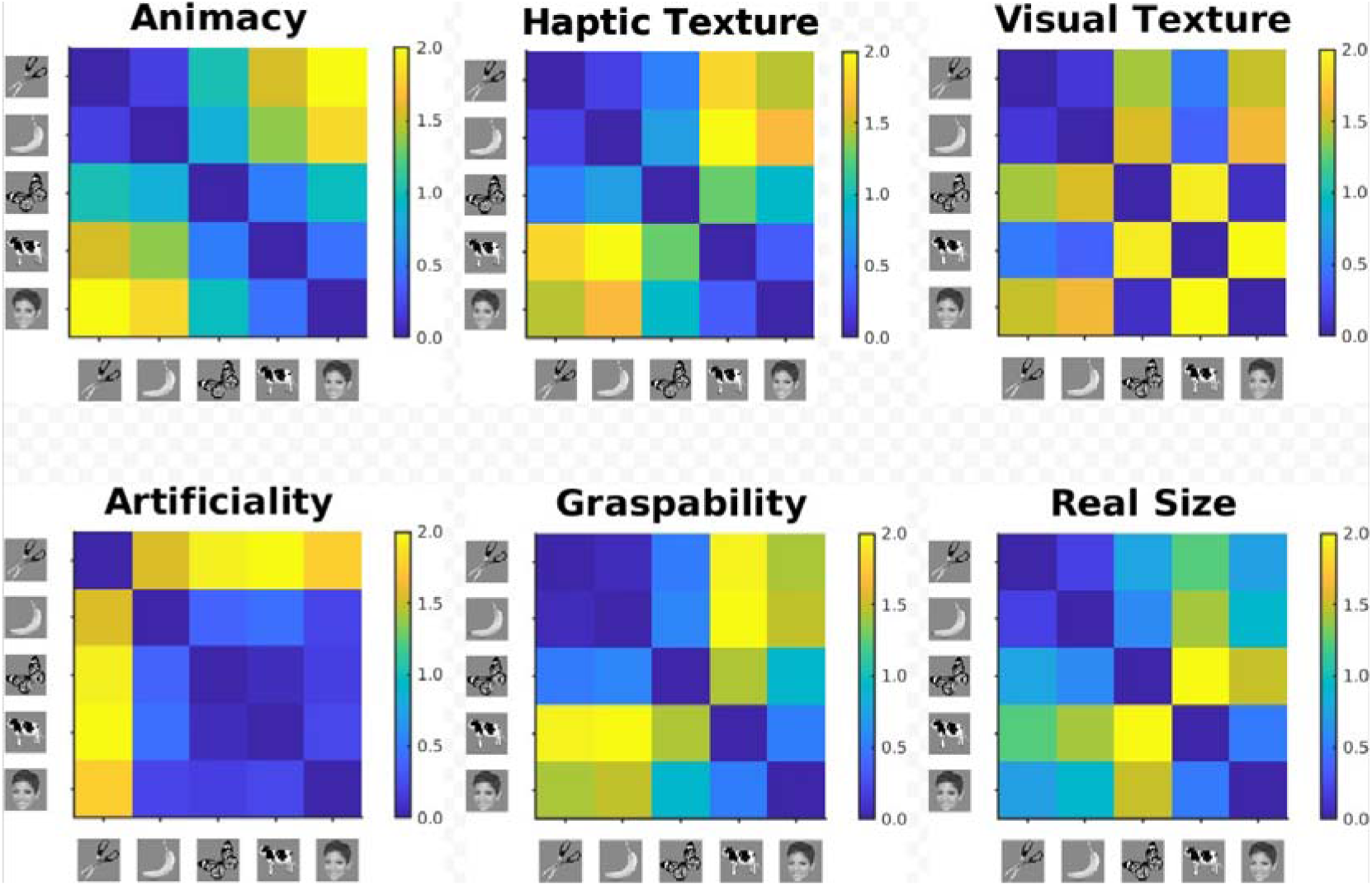
Composite behavioral RDMs. Here we present the RDMs of the six behavioral composite models computed for the RSA. Behavioral judgements were averaged within the categories, the dissimilarity between categories was calculated using their euclidean distance and lastly rearranged from 0 to 2.

The comparison between neural and behavioral RDMs was computed using the RSAtoolbox (Nili et al., 2014). Subject level spearman correlations were computed between each behavioral and neural RDM with significance corrected using FDR correction (q<0.05, for all tests within an ROI).

#### Variance Partitioning

To bring forward the conceptual understanding of how each dimension explains the neural data independently, subsequent pairwise variance partitioning analyses were conducted between the four best performing models. Variance partitioning shows us the combined explanatory power of any two models as well as demonstrating the overlap and unique contribution of each model (see Groen et al., 2018 for a similar method). With each model-pair we could see the combined variance explained (R²), the shared variance between the two models and the two unique variances explained by each model independently. The combined R² is calculated by fitting two predictors to a linear regression model explaining the neural data. In independent models, the predictors are also independently fit to a linear regression. This allows us to have three separate summary 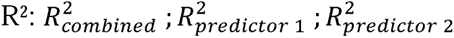. From there the shared and unique explained variances can be calculated:

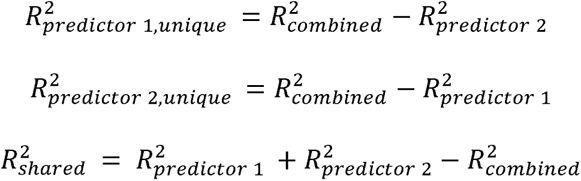

The combined R² value is the amount of variance explained by the model combination; by subtracting the individual R² from the combined value gives us the individual contribution (unique variance) of each model in the given combination; the shared variance is the sum of the two individual models subtracting the combined R².

## Results

### Is neural discriminability of the responses to the different categories of stimuli dependent on their animacy status?

In order to understand whether animacy can explain the organization of object knowledge within VTC, we tested how discriminable neural responses for each stimulus category used in our experiment are from two potential extremes of the animacy continuum – faces and tools. If animacy is the major organizing principle of VTC, then ease of discriminability, as a proxy for neural similarity along the main principle of organization within VTC, should reflect, at minimum, the animacy status of these categories. Specifically, neural discriminability should follow, potentially linearly, how animate the exemplars of these categories are. Irrespective of the definition of animacy one assumes (e.g., agency, self-motion, volition, humanness), the ranking of the categories used in our study, given animacy, should be the following: faces as the most animate, followed by animals (mammals), insects, then fruits and vegetables, and, finally, tools. Specifically, and again irrespective of the definition of animacy, because fruits and vegetables have to be considered less animate than insects – one would be hard pressed to attribute more animacy to broccoli than to a cockroach – fruits and vegetables should be easier to discriminate from faces than insects, and insects should be easier to discriminate from tools than fruits and vegetables.

To test this, we conducted multivariate pattern classification, separately in the right and left VTC, and performed binary classification between each stimulus category and the two anchor categories independently, using linear support vector machine (SVM) algorithms. Figure 4A shows the results of the SVM classifiers when individually training the decoder on all conditions versus tools (see supplementary table S2 for the statistics on pairwise comparisons). As expected, faces are highly discriminable against tools and thus are classified with very high accuracy. Importantly, and as expected given an animacy claim, animals are also highly discriminable against tools, but significantly less so than faces. However, not only are insects and fruits and vegetables hard to discriminate against tools (significantly more so than faces or animals are discriminable against tools), but there is no significant decrease in accuracy when discriminating between insects against tools and fruits & vegetables against tools. In fact, if anything, insects are (numerically; *t*s < 1) harder to discriminate against tools than fruits and vegetables are to discriminate against tools. Interestingly, Figure 4B shows similar results regarding the SVM classification of all conditions versus faces. Once again, and as expected, tools are highly discriminable against faces and thus are classified with the highest accuracy. Conversely, animals, again as expected, are the least discriminable against faces, significantly so than any other category. Importantly, results show that it is easier to discriminate fruits and vegetables from faces 1) than insects from faces (left VTC, *t* (15) = 2.07, *p* = 0.0561, FDR corrected; similar numerical trend in the right VTC), and 2) than tools from faces (right VTC, *t*(15) = 2.15, *p* = 0.0483, FDR corrected; similar numerical trend in the left VTC; see supplementary table S2). This suggests that the patterns of activation for insects are more similar to that of tools – a result that is hard to reconcile from the standpoint of the animacy continuum as the major organizing principle of VTC.

**Figure 4.**
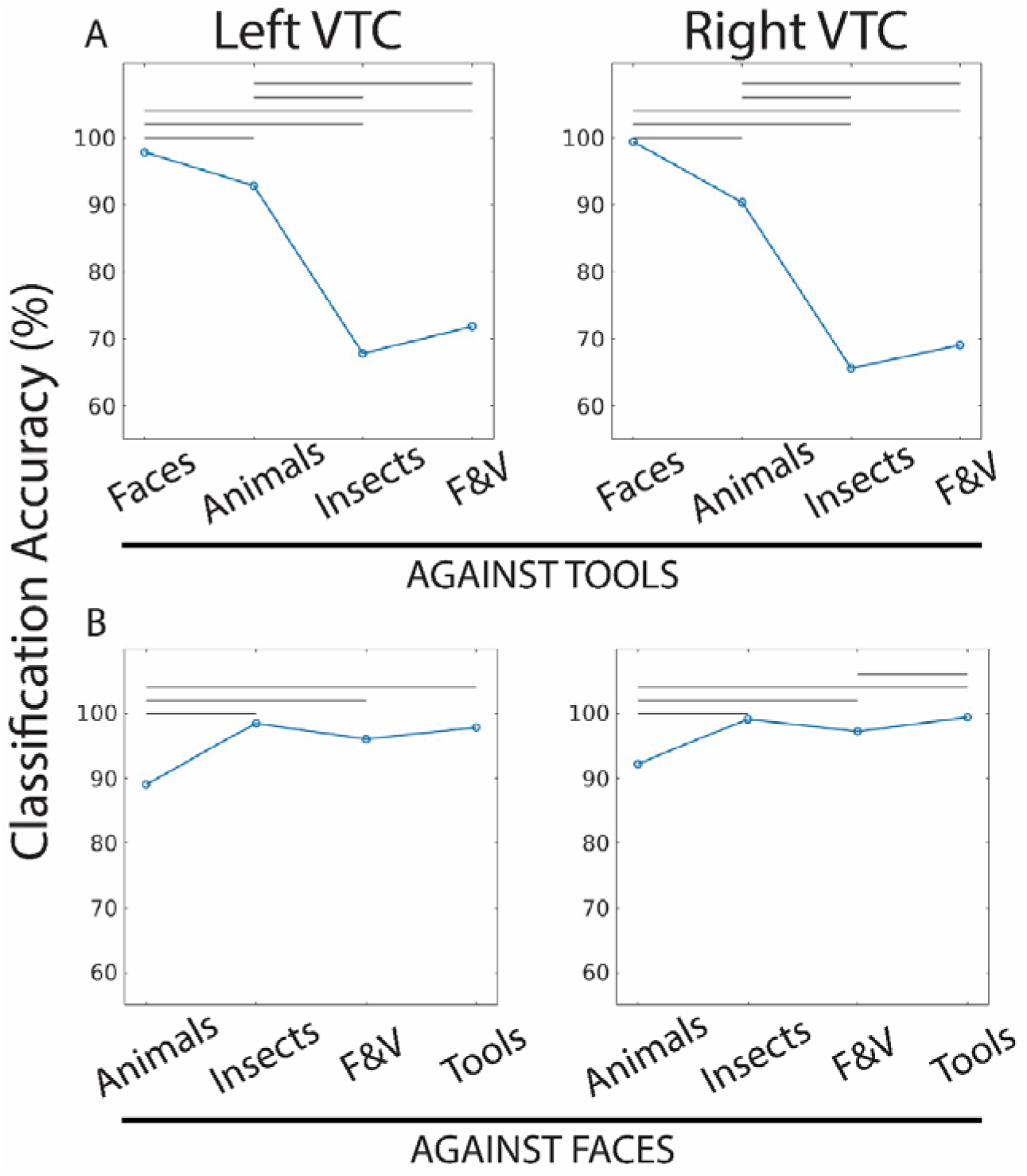
Discrimination accuracy between pairs of categories. Results from the SVM classifier for the left and right VTC. Each graph shows the accuracy of the classifier to distinguish between different pairs of stimuli categories when comparing all categories versus tools (A) and faces (B). Horizontal bars show a significant difference between two pairs at FDR corrected p<0.05. F&V: Fruits and Vegetables.

### What dimensions might best explain the representational content within the VTC?

To further characterize the organization of VTC, we focused on explaining the representational content of the VTC through a series of important object properties spanning shape, texture, action, animacy and ontological characteristics of the stimuli used herein. To do so, we used RSA (Kriegeskorte et al., 2008) to explore which stimuli properties may play a role at explaining the functional organization of VTC. Behavioral and neural RDMs were compared using FDR-corrected Spearman correlations. Figure 5 shows the similarity in representational content between the different behavioral-model RDMs and the neural RDMs for the left and right VTC. A major finding here is that haptic texture and grasping properties supersede numerically (and statistically in the right VTC; FDR-corrected, *p* < 0.001 for all measures) the ability of the animacy model (and real size model) to explain the representational content within the VTC. Importantly, the animacy model and the real size model also fare relatively well in explaining representational content within VTC, whereas the remaining models do not seem to be able to predict variance in the neural representational similarity.

**Figure 5.**
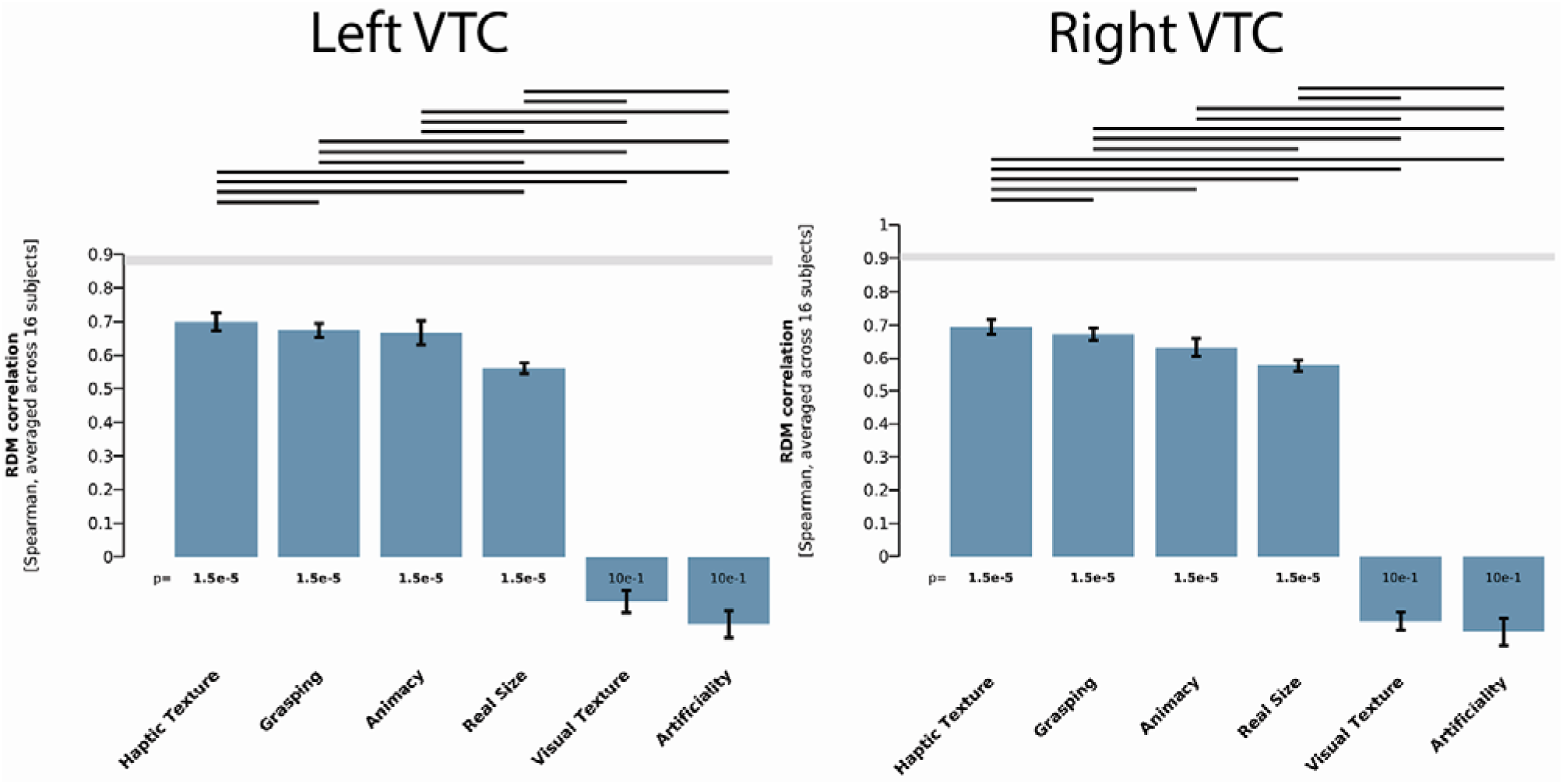
RSA results for the left and right VTC. Vertical bars show the standard error of the model correlations, and horizontal bars show significant difference between two models at an FDR corrected p<0.05. The noise ceiling in gray represents the correlation to be reached for a behavioral model to fully explain the variance in the neural activations.

One important question that arises from these results relates to potential and shared variance explained between these models. To test this, we performed pair-wise partial correlation analyses between the four models with a significant correlation to the VTC’s functional organization on the left and right VTC (Groen et al., 2018). Figure 6 shows the results for this analysis. On top of each model-pair, the variance explained jointly by the models (R²) is shown. The shared and unique variance are demonstrated as a percentage of the joint R². Firstly, given their significant correlation with the functional representations in the VTC, all models explain a high amount of variance. Secondly, haptic texture and graspability (i.e. the two best performing models in the RSA; Figure 5), explain an important amount of unique variance when pitted against animacy and real size. Furthermore, the two latter models have an extremely reduced unique variance when modeled with haptic texture and graspability, indicating that their high performance in the RSA is due mostly to shared traits with haptic texture and graspability, therefore greatly reducing their explanatory individual explanatory power. Finally, the near indistinguishable variance explained by haptic texture and graspability points to a noteworthy finding. These two models were collected using distinctly different methods: graspability was computed using a single judgement task (how easily can an object be grasped with one hand), while haptic texture was computed using an average of four different object features (smoothness, softness, furriness, and temperature), none of which could be related, at face value, to whether an object was easy to grasp or not.

**Figure 6.**
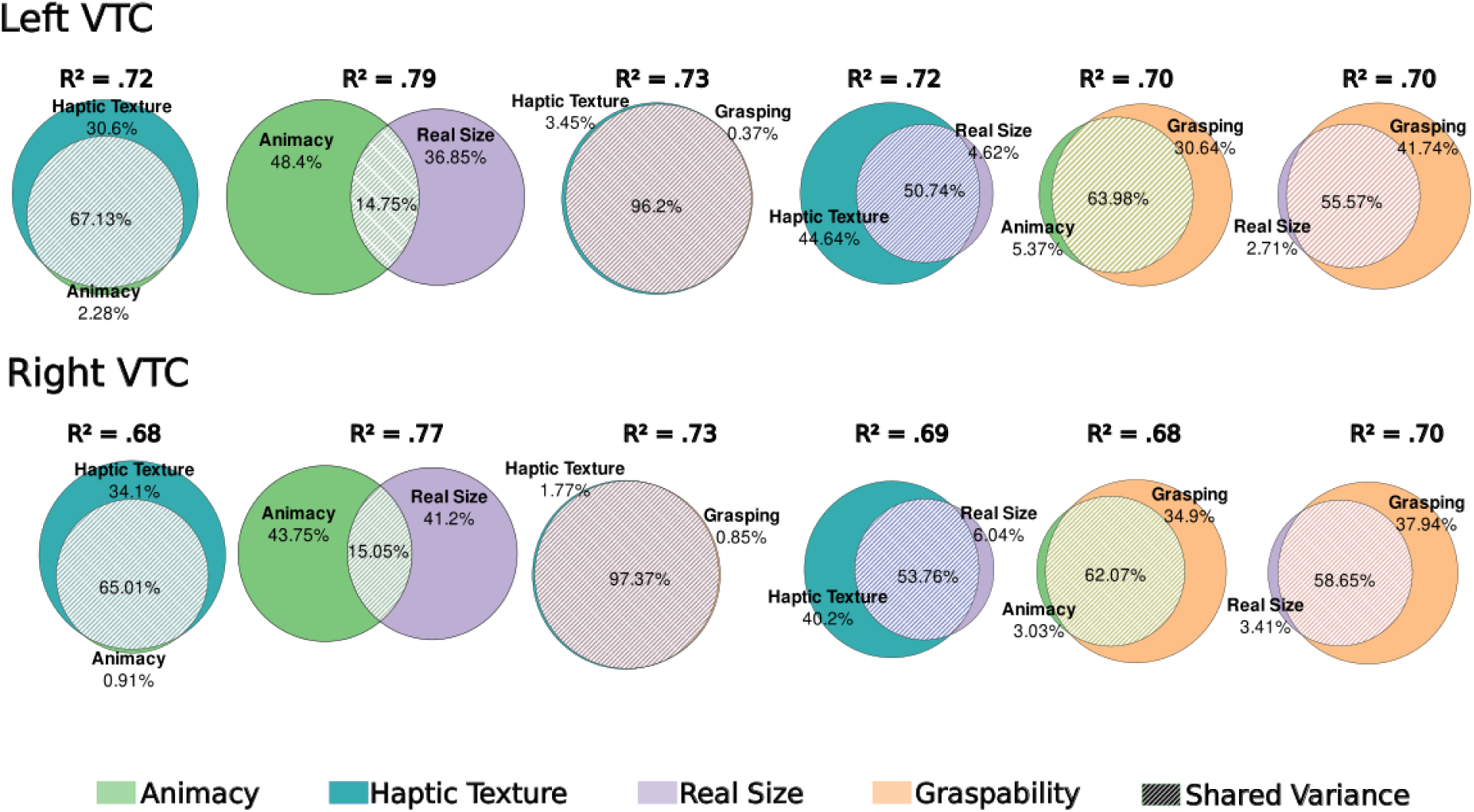
Variance partitioning between the significant behavioral models. Here we show the combined variance explained for each model pair (R²), along with the shared and unique variance of each model as a percentage of the variance explained by the target pair of dimensions.

While these two models refer to conceptually separate object features, our results show the importance of their combined integration in the functional organization of the VTC (see supplementary table S1 for the relevant category score for each model).

## Discussion

Understanding object recognition and the build-up of mental representations of the *things* that lay outside in the world is a major step in understanding the human mind. The field has focused on studying the organization of such representations in the brain as a proxy for understanding object recognition – and this has led to the proposal of different major principles of organization of representational content in the brain. One such proposal, perhaps fueled by the overall notion that the animate/inanimate status of an object is a fundamental representational property and may figure critically in the ability of spared and impaired object recognition, several authors have proposed that the major principle of organization of information in the areas that subserve object recognition – specifically in VTC – is animacy (e.g., Sha et al., 2015; Thorat et al., 2019). In particular, the organization of information on and representations of objects in VTC should follow an animacy continuum that tracks each stimulus animacy status (e.g., Connolly et al., 2012; Ritchie & Op de Beeck 2019; Ritchie et al, 2021; Sha et al., 2015; Thorat et al., 2019) in a domain-general fashion. Others have proposed dimensions such as real-size (Konkle & Caramazza, 2013; Konkle & Oliva, 2012), or action-related material and the graspable status of an object (Almeida et al., 2008, 2010, 2013, 2018; Amaral et al., 2021; Chen et al., 2017; Cortinovis et al., 2025; Garcea et al., 2018, 2019; Hussain et al., 2024; Kristensen et al., 2016; Lee et al., 2019; Mahon & Almeida, 2024; Mahon et al., 2013, 2022; Ruttorf et al., 2019; Walbrin et al., 2024; Walbrin & Almeida 2021) that may or may not relate more specifically to particular domains.

We sequentially tested these different proposals by looking at pattern discriminability and representational similarity of the neural responses elicited by different categories of stimuli. We started by testing the role of animacy and the animacy continuum in the organization of representations in VTC. We focused specifically on how two categories of objects – insects and fruits and vegetables – fare in relation to two possible anchor categories of the animacy continuum – tools and faces. While fruits and vegetables and insects are both living things, they differ in their animacy status in that fruits and vegetables are clearly less animate (if at all animate) than insects. As such, if animacy is the major principle of organization within VTC, then the patterns of activation elicited by these two categories, and their representational content within this area, should reflect their animacy status – that is, fruits and vegetables should be much more similar to tools than insects and insects should be much more similar to faces than fruits and vegetables. Our results on pattern discriminability do not seem, however, to support this contention, nor the central importance of an animacy continuum as the main organizational principle of information in VTC.

Specifically, our decoding analysis demonstrates a breakdown of the animacy continuum within VTC for the different categories tested herein. When observing how discriminable each category is from tools within the VTC, we showed a numerical increase in decoding performance from insects to fruits and vegetables (against tools), suggesting the lack of a dominant animacy dimension – that is, although insects are a more animate category than fruits and vegetables, they are not any harder to discriminate from tools (quite the contrary, at least numerically). A similar pattern emerges when comparing each category tested against face stimuli. Specifically, while fruits and vegetables are more inanimate than insects, they are not easier to discriminate against faces than insects (again, our results show a marginal statistical significance demonstrating that insects are in fact harder to discriminate against faces).

We then tested how several possible dimensions (including animacy) fared in explaining the representational content of VTC. While animacy and real-size were important dimensions, these were surpassed by haptic and action-related texture and by graspability as major explanatory dimensions of the organization of representational content in VTC – i.e., our representational similarity analysis shows that the best models overall in explaining the representational content within VTC were those related with haptic texture and the graspability status of stimuli. These models were, for the most part, not only the best predictors, but importantly, they were also better predictors of the representational content within VTC than animacy and real size. We also showed that while these dimensions – haptic texture, graspability, animacy and real-size – conjointly explain a large part of the variance in representational content within the VTC, some of them also uniquely explain a considerable part of the variance. This is especially true for haptic texture and graspability when compared dichotomously to real-size and animacy – not only do they independently explain a considerable part of the variance, but they also almost completely exhaust the contribution of real-size and animacy (see Fig. 6). Interestingly, though, haptic texture and graspability share almost all the variance explained, despite the fact that they refer to independent concepts, and were computed in a completely independent way.

These results seem to be in line with work on the multidimensional nature of object processing (Almeida et al., 2023, 2025; Amaral et al., 2022, Fernandino et al., 2021; Hebart et al., 2020; 2023; Huth et al., 2012; Valério et al., 2025, 2026) and the fact that several object-related dimensions may collectively explain the representational content within different parts of the brain. For instance, we have recently shown that dimensions related to visual (e.g., material, shape) and action (e.g., grasp type) properties of objects can explain activations within VTC and lateral temporal cortical regions (Almeida et al., 2023, 2025; Fernandino et al., 2021; Hebart et al., 2020; 2023; Huth et al., 2012; Valério et al., 2025, 2026). In fact, (at least some of) these dimensions seem to be organized in continuous topographic maps – contentopic maps (Almeida et al., 2025) – that relate to intermediate object representations bridging between early visual properties and complex object representations. Perhaps, transformations over the readouts of these contentopic maps within occipital and occipito-temporal cortex, shape and constrain the organization of the object representations downstream within more anterior VTC.

Moreover, our results also seem to be in line with early neuroimaging and neuropsychological work on the role of VTC in texture and material processing (e.g., Almeida et al., 2023; Baumgartner and Gegenfurtner, 2016; Cant et al, 2008, 2009; Cant & Goodale, 2007, 2011; Cant & Xu, 2012; Cavina-Pratesi et al., 2010; Jacobs et al., 2014; Komatsu & Goda, 2018; Prodebarac et al., 2014; Xiao & Liao, 2025). Collectively, this work has shown that the processing of surface properties, such as texture and material, is dependent on regions within the collateral sulcus. Potentially, these ventral temporal regions are coding for different kinds of haptic and visual-related properties of materials. The centrality of these kinds of computations and information on the organization of object representations in the brain has been strongly underscored recently (Mahon & Almeida, 2024). Mahon & Almeida proposed that reciprocal interactions between anterior parietal (e.g., anterior Intraparietal Sulcus – aIPS - and the Supramarginal Gyrus) and ventral and lateral temporal regions dictate local computations within these areas – for instance, an object’s material (processed within ventral temporal cortex) will strongly constrain the computation of a grasp (within aIPS) and support end-state comfort; the graspable status of an object (processed within parietal cortex) will impact the organization of representations within VTC via querying related to material and surface properties of target objects, enhancing response tuning towards graspable objects within VTC.

According to this proposal, aspects of surface properties that inform action and aspects related to the graspable status of an object should hold an important role in the organization of representations in VTC. Very in line with this, our results seem to show that surface and material properties that have an haptic aspect to it (in our study, smoothness, softness, furriness, and temperature) and that relate strongly with action and grasping (as seen by the shared variance between the haptic texture and grasping models in Figure 6) – but not those that are almost exclusively visually-detectable and may have limited effects on the grasping program to be applied to the target stimulus (in our study, texture complexity, glossiness and opaqueness) – are strong predictors of the representations within VTC. Speculatively, this most probably explains the almost complete overlap between haptic texture and graspability in VTC, strongly suggesting, as Mahon and Almeida (2024) hinted, that actionable texture and material properties – i.e., those properties that have an impact in programing grasps and interactions with target stimulus – are an important principle of organization of ventral temporal cortex.

In fact, this multidimensionality of object processing and the connectivity constraints that have been shown to be at play within VTC (Almeida et al., 2013, 2023, 2025, Amaral et al., 2021, 2022; Bouhali et al., 2014; Bracci & Peelen, 2013; Chen et al., 2017; Fernandino et al., 2021; Garcea et al., 2018, 2019; Hebart et al., 2020; 2023; Hutchinson et al., 2014; Hutchinson & Gallivan, 2018; Huth et al., 2012; Kanwisher, 2016; Kristensen et al., 2018; Lee et al., 2019; Mahon, 2022; Mahon et al., 2013; Mahon & Almeida, 2024; Mahon & Caramazza, 2011; Martin, 2016; Osher et al., 2016; Riesenhuber, 2007; Ruttorf et al., 2019; Saygin et al., 2016; Valério et al., 2025, 2026; Walbrin & Almeida, 2021), along with domain-specific pressures for particular computations that facilitate our interactions with the world, shaped the way in which mental representations within VTC are organized. Following previous proposals (e.g., the distributed domain-specific account; Mahon & Caramazza, 2011), domain-specific computations supporting social, navigation, action, and feeding decisions are potential first order principles of the organization of mental representations in VTC (and the rest of the brain), whose topography is defined through computational contingencies imposed from neural nodes within a domain-specific network external from VTC (e.g., Almeida et al., 2013; Amaral et al., 2022; Bouhali et al., 2014; Bracci & Peelen, 2013; Chen et al., 2017; Garcea et al., 2018, 2019; Hutchinson et al., 2014; Hutchinson & Gallivan, 2018 Kanwisher, 2016; Kristensen et al., 2018; Lee et al., 2019; Mahon, 2022; Mahon & Caramazza, 2011; Mahon et al., 2013; Osher et al., 2016; Riesenhuber, 2007; Ruttorf et al., 2019; Saygin et al., 2016; Stevens et al., 2015; Walbrin et al, 2024; Walbrin & Almeida, 2021). This first-order principle is then populated by multiple dimensions of interest (see Almeida et al., 2025 for a possible implementation) that satisfy specific queries and informational demands for the computations at play. For instance, readouts from contentopic maps over dimensions like grasp type, elongation, presence of metal, among others (Almeida et al., 2023; 2025) that relate strongly with particular domain-specific computations (e.g., action towards manipulable objects) will be aggregated dynamically and lead to the kinds of composite higher-level maps present within the VTC (Almeida et al., 2025). Domain-specific responses would thus arise as a manifestation of a multiple set of dimensions being satisfied and requested in order to fulfill specific computations – whereas traversing single dimensions would lead to weak or no selectivity (Op de Beeck, Haushofer, et al., 2008), evoking the “right” pattern of dimensions for a particular domain-specific computation to be maximally successful would lead to the specific response preferences observed.

One further aspect that merits some discussion relates to the overt difference between our results and those of Sha and Colleagues (Sha et al., 2015; see also: Connolly et al., 2012; Proklova et al., 2016; Ritchie et al., 2021; Ritchie & Op de Beeck 2019; Thorat et al., 2019) The central argument for an animacy continuum as a major principle of organization in Sha and colleagues resided on their principal component analysis, and specifically on the fact that the loadings on their first component for each of the basic objects used seemed to follow, according to the authors, a continuum in animacy status. In their experiment, Sha and Collaborators focused on the animal kingdom plus two tool items. That is, animacy was dependent also on a living/non-living distinction, rather than solely on animacy per se. It is important to note, however, that even with that caveat, a closer look to the loadings of the first component of their PCA solution by object type seems to show a rather clear divide between mammals (and perhaps even only primates and cat – an animal many of us consistently anthropomorphize) and other animals. In fact, and overall not unlike our results for insects, the loadings for clownfish, ray, ladybug and lobster are a lot more similar to those for hammer and key (the two inanimate and non-living objects used) than to those for birds, mammals and primates. Importantly, the inclusion of fruits & vegetables in our experiment allowed us to control for the living and nonliving divide, as well as further focus on the lower end of the animacy spectrum.

It is important to stress, though, that we are not suggesting that animacy is not an important aspect of the representations of an object – in fact our own RSA data suggests that animacy is, nonetheless, an important dimension in dictating the organization of conceptual knowledge within VTC. In fact, there are clear examples of the importance of an animate vs. inanimate distinction (Caramazza & Shelton, 1998; Konkle & Caramazza 2013; Kriegeskorte et al., 2008; Proklova et al., 2016; Wiggett et al., 2009) when it comes to understanding the nature of mental representations and our conceptual understanding of the *things* out in the world. Our data, however, seems to be at odds with any strong argument for animacy (and especially a continuous understanding of such a dimension) as the primary or major principle of organization of representations in VTC as proposed previously.

Another aspect in which our experiment differs from some of those that have typically addressed the animacy continuum (e.g., Sha and colleagues; see also: Kriegeskorte et al., 2008; Thorat et al., 2019) relates to the level of representation used in the experiments. Particularly, we focus on superordinate categories (e.g., insects), whereas others have focused on basic level categories (e.g., cat). Thus, the neural responses we are testing are elicited globally by a series of basic category stimuli (e.g., bee) that belong to the same superordinate category (e.g., insects). While this choice precludes us from making claims about individual, basic level, items, it nevertheless still allows us to test the hypothesis we set out to test in the beginning. That is, collectively or individually, insects or a bee have to be considered more animate (and potentially in no subtle way) than fruits and vegetables or broccoli. The same logic applies to the behavioral models created, and in fact, an inspection of the distribution and standard deviations of the responses per basic level item indicate that there is more within superordinate category disparity - suggesting that indeed one can aggregate these results by superordinate category.

Overall, we show that object representations within VTC are organized in a multidimensional fashion that potentially relates to domain-specific computations, rather than one continuous, fully distributed, overarching dimension such as animacy. For instance, actionable texture and graspable status of an object are one major organizing principle as they are central for a domain-specific architecture focused on computing action for our seamless interaction with the object out there in our environment. Thus, the organization of VTC will reflect the domain-specific computational demands potentially implemented via connectivity constraints imposed by areas outside of VTC that belong to particular domain-specific networks working on fundamental computations such as preparing actions towards graspable objects, scrutinizing social cues for interactions with conspecifics, mapping landmarks for navigation, or focusing on edibility and energy content of potential foods.

## Acknowledgements

This work was supported by the European Research Council (ERC) under the European Union’s Horizon 2020 Research and Innovation programme [Starting Grant 802553 “ContentMAP’’] to JA; and the European Research Executive Agency Widening programme under the European Union’s Horizon [Europe Grant 101087584 “CogBooster”] to JA. FB was supported by Fundação para a Ciência e Tecnologia (CEECIND/03661/2017).

