## Supplementary material for "Grasping at the organization of object knowledge: testing different object-related dimensions as organizational principles of ventral temporal cortex": ST1

¥ Corresponding author:

Jorge Almeida

Proaction Lab,

**Supplementary Table S1: Behavioral scores for all the behavioral models used for each category.**

|  | Animacy  (-1 least animate, 1 most animate) | Haptic Texture | Visual Texture | Artificiality  (-1 most artificial, 1 least artificial) | Graspability  (-1 least graspable, 1 most graspable) | Real Size  (-1 smallest, 1 largest) |
| --- | --- | --- | --- | --- | --- | --- |
| Faces | 0.67 | -0.01 | -0.14 | 0.47 | -0.21 | 0.15 |
| Animals | 0.31 | 0.07 | 0.27 | 0.65 | -0.45 | 0.40 |
| Insects | -0.09 | -0.21 | -0.01 | 0.61 | 0.23 | -0.60 |
| Fruits and Vegetables | -0.74 | -0.37 | -0.27 | 0.31 | 0.48 | -0.31 |
| Tools | -0.87 | -0.33 | -0.31 | -0.87 | 0.45 | -0.21 |

**Supplementary Table S2: Classification pairwise significance testing.** A) for all categories versus Tools; and B) for all categories versus Faces. Highlighted in green are the pairs with a significantly different SVM performance (FDR corrected).

**A**

***Left VTC***

| **Vs Tools** | Fruits and Vegetables | Insects | Animals | Faces |
| --- | --- | --- | --- | --- |
| Fruits and Vegetables |  |  |  |  |
| Insects | t(15) = -0.717, p = 0.4846 |  |  |  |
| Animals | t(15) = 7.730, p = 0.0000 | t(15) = 5.906, p = 0.0000 |  |  |
| Faces | t(15) = 9.442, p = 0.0000 | t(15) = 7.589, p = 0.0000 | t(15) = 3.873, p = 0.0015 |  |

***Right VTC***

| **Vs Tools** | Fruits and Vegetables | Insects | Animals | Faces |
| --- | --- | --- | --- | --- |
| Fruits and Vegetables |  |  |  |  |
| Insects | t(15) = -0.871, p = 0.3973 |  |  |  |
| Animals | t(15) = 4.815, p = 0.0002 | t(15) = 8.021, p = 0.0000 |  |  |
| Faces | t(15) = 7.101, p = 0.0000 | t(15) = 9.207, p = 0.0000 | t(15) = 4.035, p = 0.0011 |  |

**B**

***Left VTC***

| **Vs Faces** | Tools | Fruits and Vegetables | Insects | Animals |
| --- | --- | --- | --- | --- |
| Tools |  |  |  |  |
| Fruits and Vegetables | t(15) = -1.379, p = 0.1881 |  |  |  |
| Insects | t(15) = 1.000, p = 0.3332 | t(15) = 2.070, p = 0.0561 |  |  |
| Animals | t(15) = -3.530, p = 0.0030 | t(15) = -2.248, p = 0.0400 | t(15) = -3.822, p = 0.0017 |  |

***Right VTC***

| **Vs Faces** | Tools | Fruits and Vegetables | Insects | Animals |
| --- | --- | --- | --- | --- |
| Tools |  |  |  |  |
| Fruits and Vegetables | t(15) = -2.150, p = 0.0483 |  |  |  |
| Insects | t(15) = -0.565, p = 0.5805 | t(15) = 1.695, p = 0.1108 |  |  |
| Animals | t(15) = -3.715, p = 0.0021 | t(15) = -3.303, p = 0.0048 | t(15) = -3.467, p = 0.0034 |  |
